# A Deep Learning Approach to Detecting Temporal Characteristics of Cortical Regions

**DOI:** 10.1101/2023.08.16.553638

**Authors:** Ryosuke Negi, Akito Yoshida, Masaru Kuwabara, Ryota Kanai

## Abstract

One view of the neocortical architecture is that every region functions based on a universal computational principle. Contrary to this, we postulated that each cortical region has its own specific algorithm and functional properties. This idea led us to hypothesize that unique temporal patterns should be associated with each region, with the functional commonalities and variances among regions reflecting in the temporal structure of their neural signals. To investigate these hypotheses, we employed deep learning to predict electrodes locations in the macaque brain using single-channel ECoG signals. To do this, we first divided the brain into seven regions based on anatomical landmarks, and trained a deep learning model to predict the electrode location from the ECoG signals. Remarkably, the model achieved an average accuracy of 33.6%, significantly above the chance level of 14.3%. All seven regions exhibited above-chance prediction accuracy. The model’s feature vectors identified two main clusters: one including higher visual areas and temporal cortex, and another encompassing the remaining other regions.These results bolster the argument for unique regional dynamics within the cortex, highlighting the diverse functional specializations present across cortical areas.

## Introduction

While the mammalian neocortex has been thought to be composed of repetition of a canonical microcircuit [1], the diversity of the cytoarchitecture across cortical regions suggest functional specialization. For instance,the absolute receptor concentration throughout the cortical depth is different across the cortex [2],neuron density and the ratio of neurons to nonneuroncells (mainly glia) varied greatly across cortical areas and regions [3], the frontal lobe contains large stellate cells in layer 4 [4], while the parietal lobe’s layer 5 is characterized by large pyramidal cells [5]. Synaptic excitation and inhibition display systematic macroscopic gradients across the entire cortex [6]. Since electrophysiological activities result from the synchronous activity of neurons in the brain [7–10], we hypothesize that cytoarchitectural diversity is also manifested in electrophysiological properties of neuronal signals, such as electrocorticalgraphy (ECoG).

In previous research, it has been reported that different brain regions exhibit distinct electrophysiological characteristics. For Example, the spiking regularity of neurons increases as information flows from the input to the output regions of the brain [11]. Furthermore, studies utilizing ECoG and single-neuron recordings have revealed that the intrinsic timescales, derived from the autocorrelation of neural activity, become progressively longer as the cortical hierarchy goes higher. This phenomenon has been observed in both local field potentials (LFP) captured by ECoG and the spiking patterns of individual neurons [12]. Notably, distinct variations in spike patterns have been identified across different brain regions, including V1, V2, MT, vlPFC, PMd, and SNr [13].Besides, it has been investigated that there are cortical functional heteroginity [14–16]. Besides, there are gradients in cortical thickness across the cortex [17–19]. Notably, the dominant peak frequency in a brain area significantly decreases along the posterior-anterior axis, aligning with the global hierarchy from early sensory to higher-order areas. This spatial gradient of peak frequency is inversely correlated with cortical thickness, which serves as an indicator of the cortical hierarchical level [20].

In this study, we aimed to determine whether the functional heterogeneities of cortical regions are reflected in the temporal patterns of the local electrophysiological signals as measured by ECoG. To this end, we trained a deep learning model on ECoG data to identify similarities and differences between brain regions and extracted latent ECoG features. If cytoarchitectural diversity is reflected in each region’s ECoG, we expect to observe distinct temporal patterns. We hypothesize that the model can learn these patterns and predict the corresponding cortical region. Additionally, we predicted that regions with similar cytoarchitecture would display similar latent ECoG features, while regions with distinct cytoarchitecture would exhibit more divergent ECoG characteristics. We divided the cortical regions into seven areas based on the atlas (see Materials and Methods for details), and used the feature vectors obtained from the trained deep learning models to analyze the similarities among those areas.

## Materials and Methods

### Experiment Condition of the datasets

In this study, we used ECoG signals measured by electrodes covering most of the lateral cortex in four macaque monkeys during awake conditions from public database. [21, 22]. ECoG singnals were recorded from most of the lateral cortex in four macaques during awake (eyes-opened, eyes-closed), anesthetic and sleeping conditions. During the recordings, monkeys were seated in a primate chair with both arms and head movement restricted. The awake condition had eyes-opened and eyes-closed conditions. In eyes-opened condition, the monkey’s eyes were open and the monkey sat calmly. In eyes-closed condition, the monkey’s eyes were covered by an eye-mask to refrain from evoking visual response. We used ECoG data from the eye-opened condition. The sampling rate was 1kHz.

To prevent sample imbalances from different regions in the process of prediction by machine learning, we standardized the number of electrodes across the seven regions for each animal, adjusting to match the region with the fewest electrodes. During this modification, electrodes near the boundaries of cortical regions were removed to eliminate potential ambiguity regarding their regional affiliation. The following table1 shows the number of electrodes for each region, and table2 shows the length of the data for each electrode for each individual.

**Table 1.** Number of channels in each region.

| Monkey | LPFC | PMC | MSC | PC | TC | HVC | LVC |
| --- | --- | --- | --- | --- | --- | --- | --- |
| M1 | 12 | 12 | 12 | 12 | 12 | 12 | 12 |
| M2 | 11 | 11 | 11 | 11 | 11 | 11 | 11 |
| M3 | 14 | 14 | 14 | 14 | 14 | 14 | 14 |
| M4 | 12 | 12 | 12 | 12 | 12 | 12 | 12 |

LPFC - Lateral Prefrontal Cortex, PMC - Premotor Cortex, MSC - Motor and Somatosensory Cortex, PC - Parietal Cortex, TC - Temporal Cortex, HVC - Higher Visual Cortex, LVC - Lower Visual Cortex

**Table 2.** Length of data.

| Monkey | Time (min) |
| --- | --- |
| M1 | 15 |
| M2 | 15 |
| M3 | 20 |
| M4 | 20 |

### Deep Learning Model

In this study, we used a transformer-based model called TERA [23] known for its high accuracy in time-series analysis. While there are various analysis methods available for ECoG [24–27], our research employs this transformer-based model due to its proven effectiveness in handling temporal data. Originally developed for speaker identification in speech recognition research, TERA shares a similar objective, as we aimed to identify the “speaker” of cortical signals from time series data. TERA, an acronym for”Transformer Encoder Representations from Alteration”, utilizes the encoder part of the transformer architecture. In this study, we trained TERA from scratch to predict which electrode corresponds to which brain region using only single channel ECoG signals.

TERA acquires the latent features of waveform data in a self-supervised manner by reconstructing the original spectrogram through a one-to-one correspondence between the input data converted into a spectrogram and the data partially masked from the spectrogram. The following is the method used in this study to train TERA to acquire the latent space of ECoG and the corresponding feature vectors.(Figure1)

**Figure 1.**
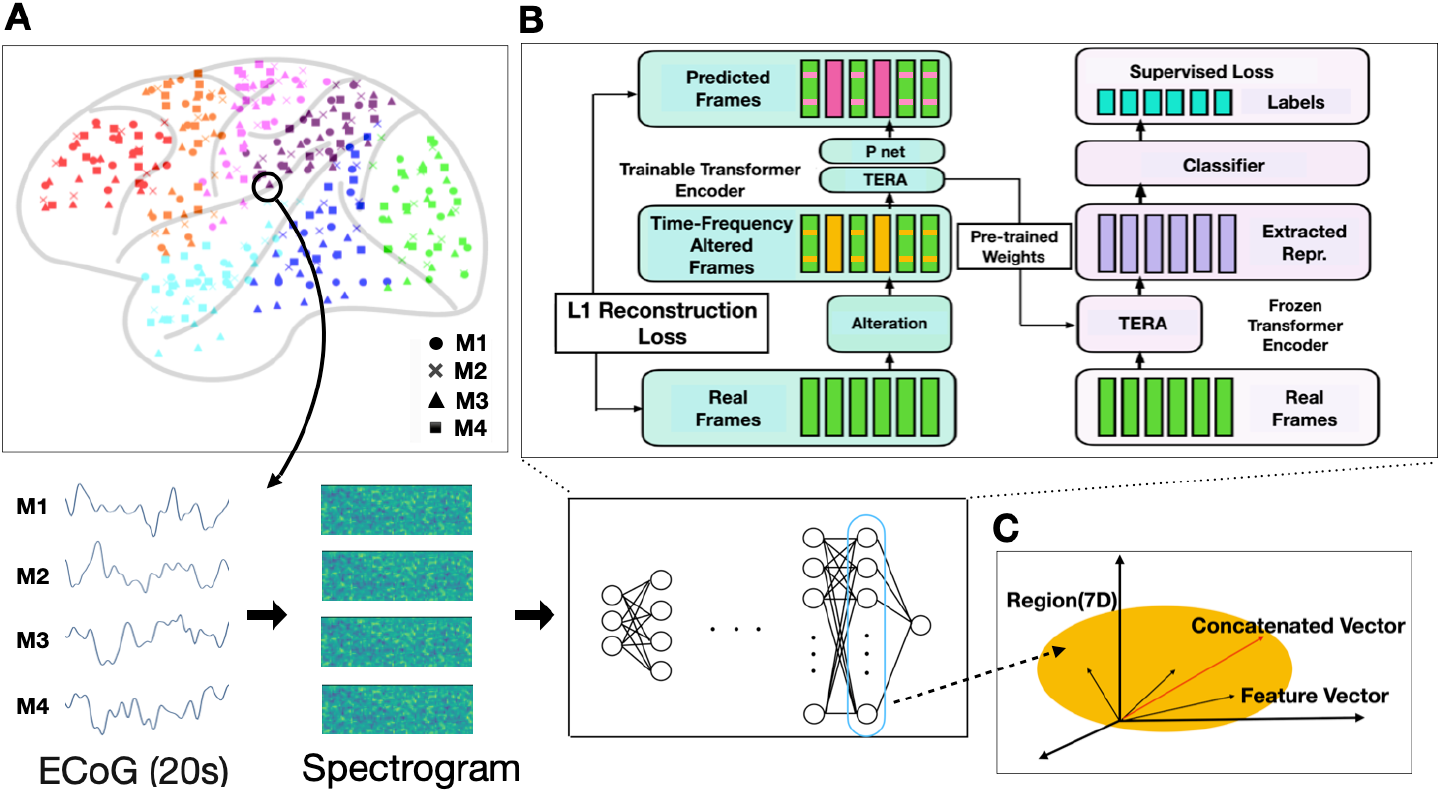
The panels in this figure show the preprocessing of ECoG data and the model trained to predict the region where the electrode belonged. (A) The cortex was divided into seven regions based on the atlas. ECoGs were cut every 20 seconds from electrodes belonging to each region and transformed into spectrograms after preprocessing. Four macaque data were used to generalize the model. (B) Model architecture. The model is trained to predict the region which input spectrogram belongs to. The model is divided into two parts, Pre-training and Downstream, with Pre-training on the left side of the architecture and Downstream on the right side. In pretraining, first, we masked the contiguous segments to zero in spectrogram along both time and frequency axis. Second, we trained the model by reconstruction the masked spectrogram. In Downstream, we fine-tuned the model so that the model predict the region the input spectrogram belongs to by supervised learning. (C) After training the model, we used the test data to extract the activation patterns of the layer before the final output layer for each input, and used those patterns as feature vectors. For each region, the feature vector was concatenated to create a concatenate vector for that region.

1. Electrode labeling according to an atlas First, the electrodes were labelled according to which region they were attached to as designated by an atlas [28] of the cerebral cortex. The placement of electrodes is depicted in Figure1.A. As described earlier, we standardized the number of electrodes across the seven regions for each animal, adjusting to match the region with the fewest electrodes 1. For each electrode, ECoG data was collected in distinct, non-overlapping segments, with each segment spanning 20 seconds. After that, the data, which was originally sampled at 1 kHz, was downsampled to 250 Hz.
2. Preprocessing Subsequently, we implemented the Common Average Reference method by averaging all channels and then subtracting the resultant signal from each individual channel. As outlined in the original TERA paper by [23], we converted the data into a log Mel spectrogram with a dimension of 80. Feature extraction was executed using a window of 25 ms and a stride of 10 ms. Following this, we applied CMVN (cepstral mean and variance normalization) to the extracted features.
3. Training the deep learning model In the final stage, we utilized the preprocessed spectrograms for model training, which consisted of two phases: Pre-training and Downstream. During the pre-training phase, the spectrograms were masked along both time and frequency dimensions. The model then learned latent spaces by attempting to reconstruct these masked spectrograms. The L1 Loss served as the loss function for this reconstruction phase.For the Downstream phase, we added a classifier to the final layer of the model using the parameters of the pre-trained model. The model was trained by supervised learning with correct labels of the regions corresponding to the input ECoG data. The model was trained to predict the electrode location from the input spectrogram. (Figure1.B) We used 80% of the data for training and 20% for the test. The same training data were used both for pre-training and the downstream task, and the test data for validation were carefully separated from this. In pre-training, we used all the training data from all the four macaques, whereas the downstream fine-tuning was done for each macaque separately. We used the data of four macaques in order to generalize the model. To discern which aspects of the spectrograms contribute the prediction of ECoG electrodes, we took two approaches. First, to investigate to what extent frequency profile contains the information about the electrode location, we tested how prediction accuracy degrades by averaging the power over time prior to model input. Second, to gauge the contribution of temporal patterns, we randomized shuffled all the time points in a randomized order before feeding them to the model. These analyses are expected to provide a hint as to which aspects of the spectrograms contain the characteristics of brain regions that contributed to successful predictions. In order to quantify to what extent the model managed to predict the region of the target ECoG electrode, we calculated F1 value for each region. F1 value is the harmonic mean of two other metrics, precision and recall. The formula is as follows;

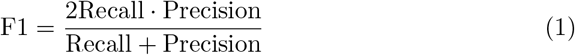

Precision represents the proportion of correct predictions made by the model among all the predictions it made. Recall, on the other hand, represents the proportion of correct predictions made by the model among all the actual positive cases.
4. Saliency Map To investigate which parts of the spectrogram the model relied on, we computed saliency maps. Saliency maps are widely used in deep learning as a technique to understand the importance or relevance of different regions within an input image or data. They provide a visual representation that highlights the most significant areas or features that contribute to a model’s prediction. In this study, saliency maps were generated using gradient backpropagation. These methods exploit the gradients of the model’s output with respect to the input features to determine the importance of each feature. The gradients were computed by backpropagating the model’s output through its layers, attributing relevance to the input features based on their impact on the final prediction. This process allows the identification of the most influential features that contribute positively or negatively to the model’s decision. Once the gradients were obtained, they were visualized as a saliency map, which is a heatmap overlayed on the input image. The heatmap assigned different intensities of colors to different regions, with the most salient areas represented by brighter colors or higher intensities. These salient regions correspond to the areas that strongly influence the model’s prediction.
5. Extraction of latent feature After training the model, the output of the one before the last layer of the model was extracted for each input. The layer is circled in the Figure1.B. This output essentially serves as a feature vector within the resultant latent space. Based on this feature vector, we gauged the similarity amongst regions. That is, small distances between feature vectors indicated higher similarities (Figure1.C). The feature vectors were added together for each region, and those added vectors were used as the concatenated vectors in that region. We calculated and compared the distances of concatenated vectors for each of the seven regions, under the assumption that the distance between concatenated vectors across regions reflects inter-region similarity.

### SVM

To evaluate whether the transformer-based model had any advantage over other simpler methods, we obtained a simple baseline performance using Support Vector Machine (SVM). We used the same spectrograms as input for training a linear SVM. The data preprocessing technique used for the TERA model was applied to preprocess the spectrogram in the same way. This involved performing the common average referencing (CAR) and cepstral mean and variance normalization (CMVM) subtraction as preprocessing steps. For each individual, an SVM was trained using the labeled electrode regions as the target labels, with the preprocessed data serving as input. After the preprocessing, the spectrograms were transformed into one-dimensional vectors of (feature length)×(time length). The training data consisted of 80% of the overall data, while the remaining 20% was used as the test data. The data were standardized before training the model.

## Results

### Prediction accuracy

The prediction accuracy as indexed by the F value per region for each individual is shown in Figure2.A along with the average across four animals per region. As shown, the transformer-based architecture successfully classified the region based on a single channel ECoG signal over 20 seconds for almost all region for all the four animals. The prediction accuracy of the HVC and LVC regions were relatively high, while the accuracy for the PMC region was lower than in the other regions. These results support our hypothesis that the temporal patterns in single channel ECoG signals contain some information about the recorded region.

**Figure 2.**
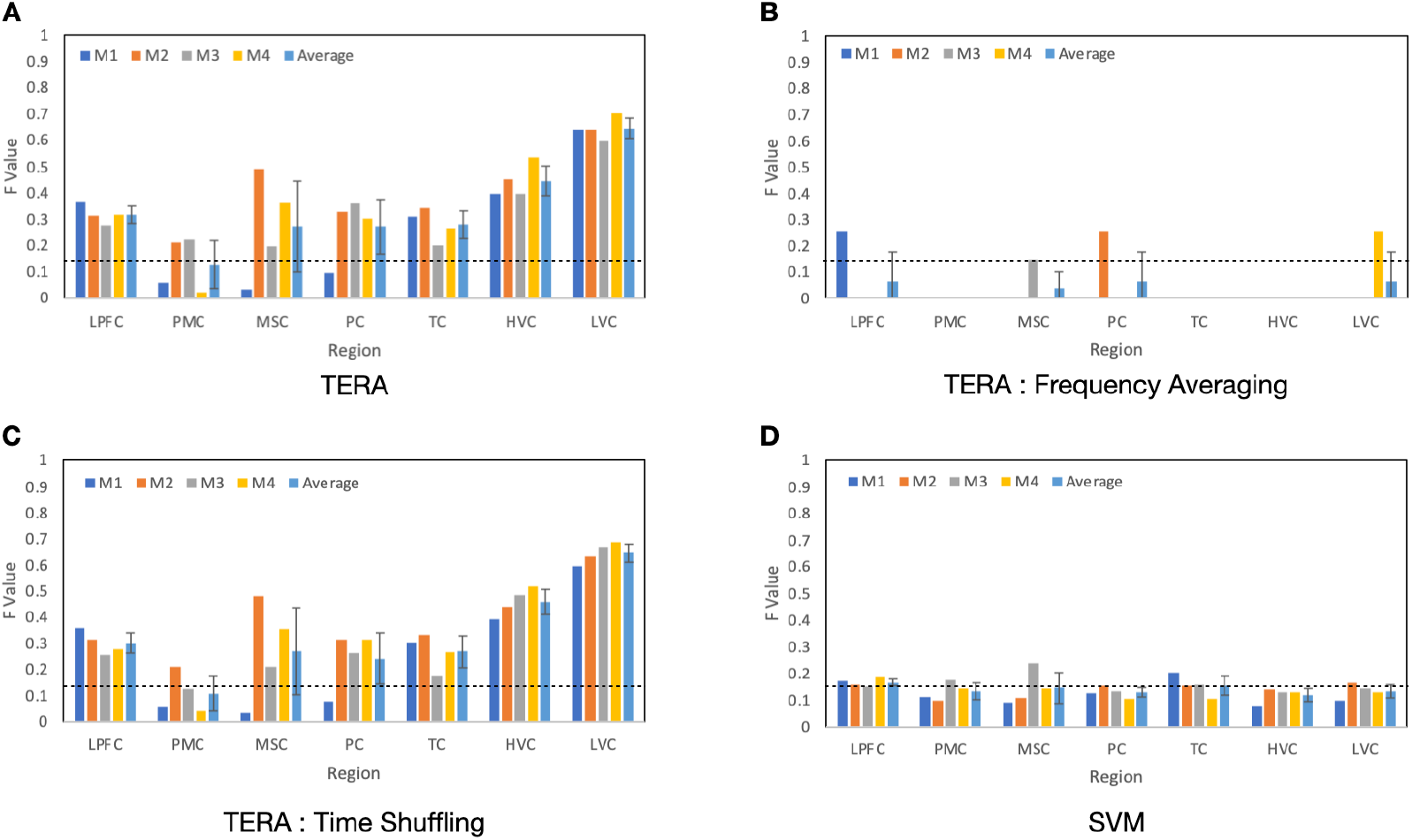
These figures show the F-values for each region in each individual when trained with models TERA and SVM. M1 to M4 correspond to each individual, and “Average” indicates the average F-values across the four individuals for each region.The dashed lines in the figures represent the chance level(1/7). (A)This figure shows the F value between individuals in each region using TERA. (B)This figure shows the F value between individuals in each region using spectrogram averaged the frequency components. (C)This figure shows the F value between individuals in each region using spectrogram, of which temporal component was shuffled. (D)This figure shows the F value between individuals in each region using SVM.

However, the specific attributes of the signals that contributed to the predictions are not clear. To address this, we hypothesized that the frequency profiles of the regions might play a role. To evaluate this, we calculated the average power for each frequency by compressing the spectrogram over its temporal dimension, and then assessed the trained model to determine any degradation in prediction accuracy. If the model relies on primarily frequency information only, it is expected that even if the temporal information is removed by averaging, prediction accuracy would remain high. On the other hand, if the model relies on the combination of both frequency and temporal information, then the averaging would degrade the accuracy.

The results of this analysis are shown in Figure2.B. Contrary to our expectation, the prediction accuracy was at chance levels. This implies that the predictions were not solely reliant on the frequency power profile. Instead, the model seems to be attuned to more intricate features within the time-frequency data.

To further understand the contributing features, we shuffled the time of the input spectrogram to see how much the model relied on temporal features. Figure2.C shows the F value between individuals in each region using spectrogram, of which temporal component was shuffled. It is observed that the F-values did not change significantly compared to Figure2.A. This suggests that the temporal structure in the spectrogram plays little role in the prediction of the region. The combined results from these two additional analyses may be somehow perplexing because powerspectrum alone is not sufficient for successful prediction, suggesting the relevance of temporal information, whereas shuffling the temporal order did not harm the performance. Possible interpretations of these results will be discussed in the discussion section.

Finally, we show the results from our baseline performance obtained with linear SVM. Figure2.D shows the F value between individuals in each region using SVM. From this figure, it appears that the F-values are close to the chance level. This suggests that when using the simple baseline model SVM, as compared to TERA, it was not able to capture the ECoG features effectively in each region.

In addition to those attempts to reveal contributing features by degrading the input data, we also applied methods to visualize which portion of the data the model paid attention to for making predictions. Figure3 visualizes the saliency map of the model using the gradient of the model. The horizontal axis is time and the vertical axis is frequency, and the Saliency Map visualizes the features that the model pays attention to. The trial average, the average of correct cases, and the average of incorrect cases were calculated for each region for each individual. The highlighted horizontal bands in those figures suggest that the model focuses mainly on frequency components. The appearance of temporal changes at a certain moments may suggest that the model may be capturing a temporal feature. On a single trial basis, we often observed temporally specific attention that appeared as brief vertical lines that overlapped with high gamma activities. Such brief events may have also made contributions to the prediction accuracy.

**Figure 3.**
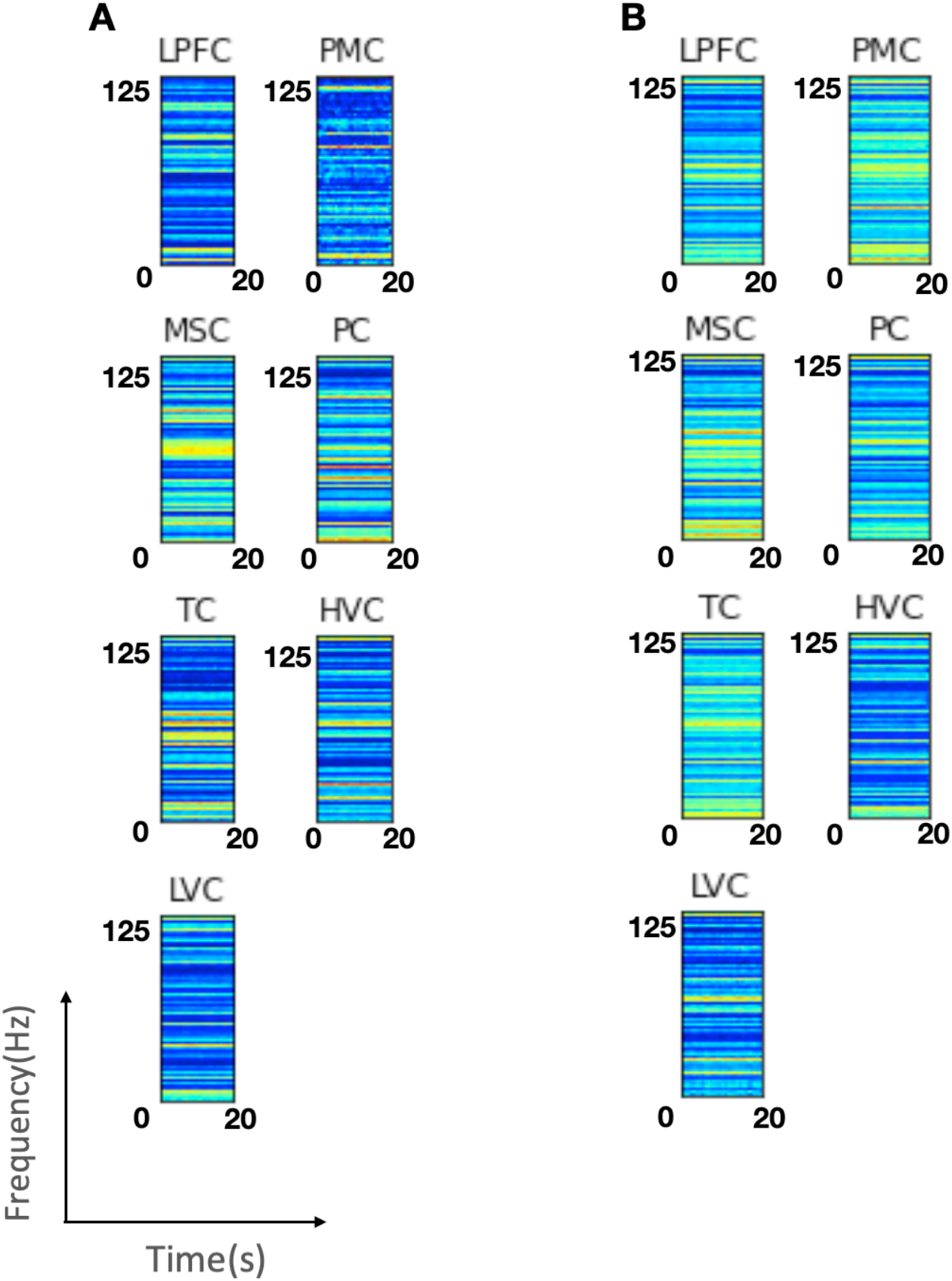
These figures represents saliency map of each regions (A)This figure represents the averaged spectrograms when the prediction was True. (B)This figure represents the averaged spectrograms when the prediction was False.

### Similarities of cortical regions

In order to assess the similarity between regions, we extracted the activation patterns from the layer just before the last layer of the trained model. This vector was used as the feature vector of the region to which the input ECoG data belonged, and this feature vector was used to examine the similarity between the regions. The similarity between regions was quantified using Cosine Similarity. For each region, the feature vectors were extracted for each 20-second segment and then concatenated to make a long vector spanning over the entire period. Similarities between regions were computed based on those concatenated vectors obtained for each region (Figure1.C) as follows. Let a and b be two representative vectors, the distance between vectors can be obtained as

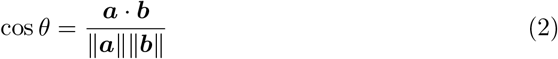

The Cosine Similarity diagram (Figure4) shows that both HVC and TC were independent of other regions and the other regions forms one cluster. Within the big cluster, MSC and PMC, and LVC and PC, respectively formed subclusters. These results suggest that ECoG signals derived from the temporal cortex exhibit more distinct characteristics compared to other regions.

**Figure 4.**
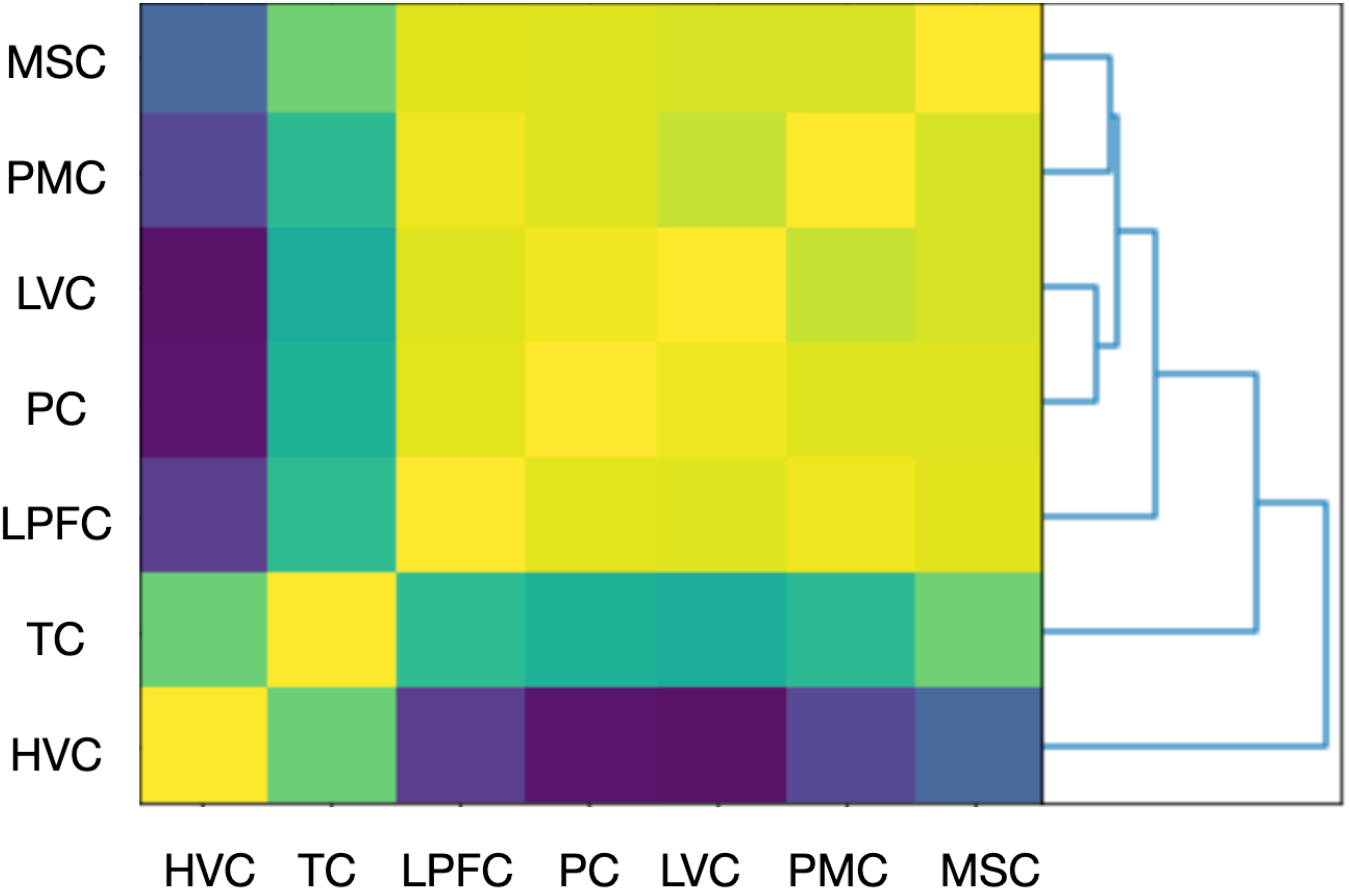
This is the Similarity matrix between the regions. Concatenated Vectors were created by adding the Feature Vectors for each region, and the distance between the concatenated Vectors for each region was calculated using cosine similarity.

## Discussion

The purpose of this study was to test whether different cortical regions exhibit distinct electrophysiological characteristics. We used ECoG signals obtained from each electrode for classifying the corresponding electrode’s region into one of seven categories, as delineated by anatomical criteria. Our anticipation was grounded in the notion that the diversity in cytoarchitecture and distinct connectivities across regions would manifest as discernible temporal patterns in their respective ECoG signals. Thus we hypothesized that deep learning models would be able to learn those distinct temporal patterns and determine which region the ECoG belonged to.

Our results showed that a deep learning model called TERA can indeed predict the region of an electrode based on the spectrogram computed from a 20 seconds of single channel ECoG signals. While this indicates that the temporal patterns of ECoG signals contain the characteristics pertinent to the target region, it was not clear what patterns actually enabled the prediction. It was perplexing that while simple frequency profiles were not sufficient for successful prediction, shuffling the spectrograms in the temporal domain did not substantially impair the prediction accuracy. The results of our saliency mapping did not provide unequivocal insights as to the underlying contributing features.

A plausible interpretation for these results is that our model identified transient events specific to each region. For instance, upon examining the saliency maps for individual time segments, we sporadically noticed fleeting highly salient moments across all frequencies coinciding with high gamma activation. These transient events coupled with other subtle attributes may have played a role in making predictions. However, we were not able to pinpoint definitive patterns that consistently served as hallmark features for regional classification. This remains a significant constraint of our present study and a more in-depth exploration is required to comprehensively grasp these underlying characteristics.

In addition to showing the possibility of predicting the region of the electrode, we explored the similarities and differences in the ECoG signatures across brain regions. To this end, we used the trained neural network models as a way to create feature vectors for each region and constructed a similarity matrix across the seven brain regions. The results shown in Figure.4 show that brain regions are split into two clusters: the HVC and TC formed one cluster while others made another. This suggests that ECoG signals in the temporal cortex are distinct from the rest of the cortex. Those regions in the temporal cortex belonged to the so-called ventral visual pathway associated with high-level vision [29–31]. Additionally, we observed subtler subclusters, denoting similarities between the early visual cortex and parietal cortex, as well as between somatomotor and premotor areas. These cortical configurations hint at a division of dynamics into the ventral pathway, dorsal pathway, and motor-related clusters.

Such demarcations might reflect diverse characteristics of the inherent neuron populations. For example, the contrasts between the ventral and dorsal pathway clusters might originate from variations in their connectivity and functions. The ventral and dorsal pathways are known for their distinct architectures, topographies, and connections [32]. Neurons in the frontal lobe are markedly more spinous than their counterparts in other lobes. [33]. Differences manifest in the spine count of the basal dendritic trees of pyramidal cells across V1, V2, and PFC, as well as in their dendritic tree sizes [34]. Moreover, distinctions in spine size, cell count, and size between the dorsal and ventral stream areas have been documented [35, 36]. According to quantitative cytoarchitectonics, it is observed that dorsal and ventral pathways are separated in the clusters [37]. Any of these variations may contribute to the unique temporal dynamics in ECoG signals that allowed for the classification of the region of origin.

In summary, our study aimed to determine whether distinct cortical areas exhibit unique electrophysiological markers. Utilizing ECoG signals and the capabilities of the TERA deep learning model, we successfully classified electrodes into one of seven regions. This classification underscores the potential variations within the cortex.

However, the specific patterns responsible for this accurate classification remain less clear, especially given the unexpected results from our frequency profile and time domain experiments. We speculated that transient events and other subtle features that might play crucial roles in predicting the region from ECoG signals. In examining similarities and differences in latent features across regions, we found evidence suggesting that ECoG dynamics could be segmented into distinct clusters. These observations might reflect underlying neuronal properties, ranging from dendritic structures to cytoarchitectural differences. Our findings have added to the existing body of knowledge on the cortex’s electrophysiological properties, and we anticipate further research will continue to clarify these complex relationships.

## Acknowledgments

This work was supported by JST, Moonshot R&D Grant Number JPMJMS2012.

